# Effect of 40Gy irradiation on the ultrastructure, biochemistry, morphology and cytology during spermatogenesis in the southern green stink bug *Nezara viridula* (Hemiptera: Pentatomidae)

**DOI:** 10.1101/171991

**Authors:** L. D. Stringer, D. P. Harland, J.E. Grant, J. Laban, D. M. Suckling

## Abstract

A study on the *Nezara viridula* male gonad cells was undertaken to compare normal development and development of insect cells after irradiation of 4^th^ instar nymphs with a dose of 40 Gray. Visually, morphologically, biochemically and cytologically the insects were not all uniformly affected by the radiation. In all aspects of development there was a range from severely affected to nearly normal. Irradiated insects appeared to move slowly and were unable to mate with non-irradiated females. The testes of the males varied from bright orange to grey in colour and all were smaller in size than non-irradiated testes. The ultrastructure of the developing sperm showed abnormalities the axonemes, the mitochrondrial derivatives, nebenkern and centrioles. Cytochemically, the main difference observed was the presence of granules heavily stained with acid phosphatase in between mitochrondrial derivatives. The chromosomes of these irradiated insects were highly fragmented. Although a few sperm in irradiated insects appeared normal no progeny were produced as insect did not mate. The sterile insect technique (SIT) requires a balance between the effective radiation dose to achieve partial or full sterility, while maintaining physical fitness. The observation that abnormalities varied from almost none to severe at 40Gy could help the development of SIT for control of *N*. *viridula* and other Pentatomidae.

## Introduction

The southern green stink bug, *Nezara viridula* (Pentatomidae) is an important cosmopolitan pest that is highly polyphagous on many crops, with preference for legumes and brassicas (Mau et al. 1967, DeWitt and Godfrey 1972, Todd 1989, Panizzi 1997, Panizzi and Mourão 1999, Esquivel and Ward 2014). *Nezara viridula* feeds on all above ground parts of the plant. Control relies heavily upon insecticides, most of which are disruptive to beneficial insects. The development of more benign and targeted methods for the control of stink bugs in general is desirable (Žunič et al. 2002, Knight and Gurr 2007). The Sterile Insect Technique (SIT) is one such strategy to achieve control of insect pests (Knipling 1955, Klassen and Curtis 2005, Paoli et al. 2014). SIT is species specific with no negative, off-target effects, and can be very efficient when inherited sterility is available, as in Lepidoptera (Carpenter et al. 2005) and potentially Hemiptera (LaChance et al. 1970). This is the result of viable but completely sterile and physically competitive individuals in the F1 generation. Relatively little research has been done on radiation-induced damage of the gonads or the potential for inherited sterility of Hemiptera. Reported studies have dealt mainly with the determination of effective sterilizing doses, and with the biotic effects of such sterilization (Ameresekere et al. 1971). Partial sterility of adult *Rhodnius prolixus* (Hemiptera: Reduviidae) and *N*. *viridula* was achieved after exposure to ionising radiation, with evidence of fertility rates returning to normal over subsequent generations (Mau et al. 1967, Maudlin 1976). In *N*. *viridula*, eggs were more radio-resistant than the adults (Mau et al. 1967, Sales 1977). As little as 20 Gy has been reported to be enough to render adult *N*. *viridula* sterile, and at lower doses of radiation, adults produced a higher proportion of non-viable eggs and had significantly lower fecundity than non-irradiated insects (Dyby and Sailer 1999, Žunič et al. 2002).

While there are numerous reports describing the structure of spermatozoa and spermatogenesis of non-irradiated hemipterans (Dallai and Afzelius 1980, Afzelius et al. 1985, Báo and de Souza 1994, Fernandes and Báo 1998, 1999, Fernandes et al. 2001, Özyurt et al. 2013), detailed effects of gamma radiation on the structure of spermatozoa and spermatogenesis of male hemipteran insects including *N*. *viridula* has not yet been reported. Fernandes and Báo (1998, 1999) described the normal ultrastructure of the spermatids and spermatozoon in *N*. *viridula* by transmission electron microscopy. The spermatids develop to form a highly differentiated cell, the spermatozoon which consist of a head containing the nucleus and acrosome, and the tail which is formed by axoneme and two mitochondrial derivatives. Several enzymes including acid phosphatase and glucose-6-phosphatase may be involved in this process. Acid phosphatase is associated with the Golgi complex, acrosome and axoneme (Anderson 1968, Bigliardi et al. 1970, Báo et al. 1988, Báo and de Souza 1994, Grab et al. 1997). In spermatids, glucose-6-phosphatase is associated with the endoplasmic reticulum and Golgi complex (Báo and de Souza 1994, Furtado and Báo 1996). It also associated with the acrosome of spermatozoa (Fernandes and Báo 1999).

As in many hemipteran insects, *N*. *viridula* possesses holocentric chromosomes with six autosomal pairs plus an XY set of sex chromosomes where XX is female and XY is male. One pair of the autosomes can be easily identified. It has been designated A1 because of it large size and can be either a ring or rod bivalent (Camacho et al. 1985, Papeschi et al. 2003). Irradiation in insects can cause chromosomal fragmentation and rearrangements (LaChance et al. 1970, LaChance and Graham 1984). More recently Žunič *et al*. (2002) and Dyby and Sailer (1999) found that low doses of radiation (5Gy and <10Gy respectively) affected the reproductive fitness, although there were cases of full ‘recovery’ when wildtype and irradiated progeny were mated.

Given that testis function is a key factor in successful development of the sterile insect technique it is important to understand what radiation damage does to the insect. For release of irradiated insects there needs to be a balance between the damaging effects of irradiation affecting sterility and the ability of the insect to survive and mate to continue to compromise the fertility of the wild population.

The aim of this study was to investigate the morphological, biochemical, cytological and ultrastructural changes of male *N*. *viridula* caused by exposure to 40Gy radiation.

## Materials and Methods

### Insects

*Nezara viridula* eggs were field collected from Auckland, New Zealand (-36.87, 174.75) and maintained in a laboratory at Lincoln. After hatching the nymphs were fed on fresh green beans *(Phaseolus vulgaris)* and raw peanuts (*Arachis hypogaea*) in an environmental chamber maintained at 25 ± 2^o^C, 50 ± 5% RH and 16:8 h (L:D) photoperiod following the method of Panizzi and Mourão (1999). Fourth instar nymphs were collected from the colony after two generations, and were used for the subsequent trials.

### Irradiation

To ensure unambiguous radiation damage to bug gonads, the fourth instar nymphs were irradiated to a dose of 40 Gy using a Theratron T-80 ^60^Co teletherapy external beam treatment unit (ESR, Christchurch). The insects, contained in Petri dishes (90 mm diameter, 15 mm deep), were placed at a distance of 50 cm from the radioactive point source. This point source geometry limited the dose gradient through the sample to 6%. A four millimetre thick piece of Perspex was added to the beam entrance side of the containers to ensure that full dose deposition to the insects occurred. The irradiated nymphs were returned to the aforementioned rearing conditions and were maintained until they moulted into the adult stage. All insects were dissected 24-48 h post final moult.

### Enzyme Cytochemistry and Transmission electron microscopy (TEM)

Non-irradiated (control) and irradiated live male *N*. *viridula* were placed in a refrigerator at 4°C for 15 min to reduce their activity prior to dissection. After placing them in chilled dissection buffer (0.1 M phosphate, 3% sucrose, pH 7), dissection was immediately performed using a scalpel and small dissection scissors to remove their head, legs, scutellum, wings, and abdominal integument to expose the viscera and locate the testes. The testes were carefully removed into the buffer and processed for enzyme cytochemistry TEM, or directly for TEM.

### Enzyme cytochemistry TEM

The testes were transferred directly to a light fixative (1% glutaraledehyde 0.1 M cacodylate buffer) for 15 minutes on a rotator. Samples were washed in buffer (0.1 M cacodylate, 5 mM CaCl2, pH 7.2) and cytochemical samples were transferred to either an acid phosphatase assay solution (7 mM cytidine-5’-monophosphate, 2 mM cerium chloride, 5% sucrose in a 0.1 M tris-maleate buffer at pH 5.0) or into a glucose-6-phosphatase assay solution (5 mM glucose-6-phosphate, 5 mM manganese chloride, 4 mM cerium chloride, 5% sucrose in 0.1 M tris-maleate buffer at pH 6.5; following (Robinson and Karnovsky 1983a, b) for 1 hour at 37°C, then washed with buffer.

### TEM

Testes from cytochemically-incubated samples (irradiated and non-irradiated) and non-incubated samples (irradiated and non-irradiated) were then processed for TEM by transferring directly to the primary fixative (4% formaldehyde, 2.5% glutaraldehyde, 0.1 M cacodylate, 5 mM CaCl_2_, 3% sucrose, pH 7.2) where they were fixed for 4 h on a rotator. Primary fixation and subsequent steps before polymerisation were carried out at room temperature. Samples were then washed in buffer (0.1 M cacodylate, 5 mM CaCl_2_, pH 7.2) and transferred to the secondary fixative (1% osmium tetroxide and 0.8% ferricyanide in 0.1 M cacodylate buffer) for 2 h on the rotator, washed in ultrapure water, and dehydrated through an acetone series (70%, 80%, 90%, 15 min each, then 100% EM-grade dry acetone twice for 20 min). Samples were infiltrated and embedded in procure 812-araldite 502 resin (50% resin/acetone, then thrice in 100% resin and polymerised for 22 h at 60°C. Sections 80–100 nm thick were cut on a Leica Ultracut UCT fitted with a Diatome 45° diamond knife onto 100 mesh formvar coated copper grids, post-stained briefly with 2% uranyl acetate then 0.02% lead citrate and viewed with a Morgagni (FEI; fei.com) transmission electron microscope (TEM) operating at 80 kV.

### Chromosome Analyses

Control and irradiated testes were dissected and fixed in 4:3:1 (chloroform: ethanol: acetic acid) for 4 hours followed by storage in 80% ethanol. Individual testes were softened in PBS with 2% trypsin freshly added, for 1-2 hours before squashing in 45% acetic acid and staining in 2% lacto-propionic orcein.

## Results

### 1. Structure

The reproductive organs of male *N*. *viridula* comprised paired testes, vasa deferentia, vesiculae seminalis, as well as accessory glands. The testes were elongate ovoid in form and lay across the body cavity (Fig 1A). Figure 1A shows the shape and colour reproductive organs from non-irradiated males, with testes covered by an orange peritoneal sheath (see Esquivel (2009) for further images). The irradiated gonads had the same position in the body cavity but were smaller in size, and variable in colour. In three of the four irradiated samples, their colour was orange, but the size of the testes significantly reduced (Fig. 1B). In one case, the testes were grey in colour, small in size and appeared deflated in shape, also vasa deferentia and the accessory glands appeared grey (Fig. 1C). Irradiated testes that were investigated with TEM were small in size, but not grey in colour *i*.*e*. similar to Fig.1B.

**Figure 1.**
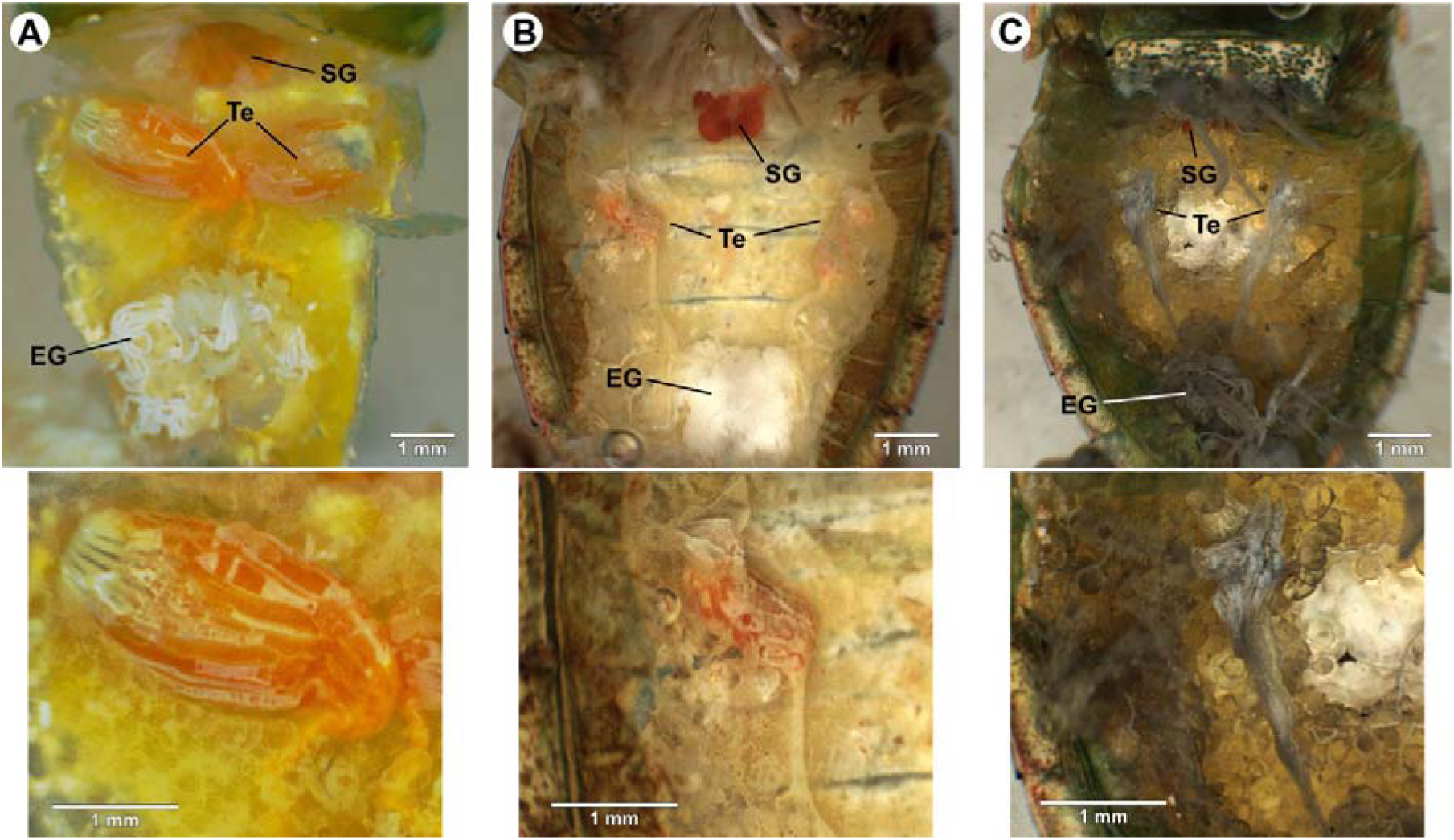
Structure of male *Nezara viridula* reproductive organs. (A) non-irradiated male; (B) 40 Gy irradiated male - similar colouring to non-irradiated male, although testes reduced in size; (C) 40 Gy irradiated male - note grey colouring of testes and other organs. Key: Te, testes; SG salivary gland; EG, ectadene gland.

In testes from non-irradiated insects, spermatogenesis proceeded in blocks of cells, within which cells were all similarly oriented and at a similar stage of development. Late spermatocyte and early spermatid stages proceeded as follows. The mitochondria aggregated close to the nucleus (Fig. 2A) and then fused together to form a single structure, the nebenkern (Fig. 2B). The nebenkern divided (Fig. 2C) and then each part elongated, taking on a vacuolated appearance (Fig. 2D) to form the mitochondrial derivatives (Fig. 2E), which eventually sat either side of almost the entire length of the axoneme of the mature sperm’s flagellum. In irradiated *N*. *viridula*, the development of the mitochondrial derivatives was less clear. Nebenkern were rare and often appeared irregular and slightly swollen (Fig. 2F). Typically nebenkern and all subsequent stages had a distinctive vacuolated appearance (Fig. 2G), about a quarter of mitochondrial derivatives having large disruptions within them, even in cells close to maturity (Fig. 2H).

**Figure 2.**
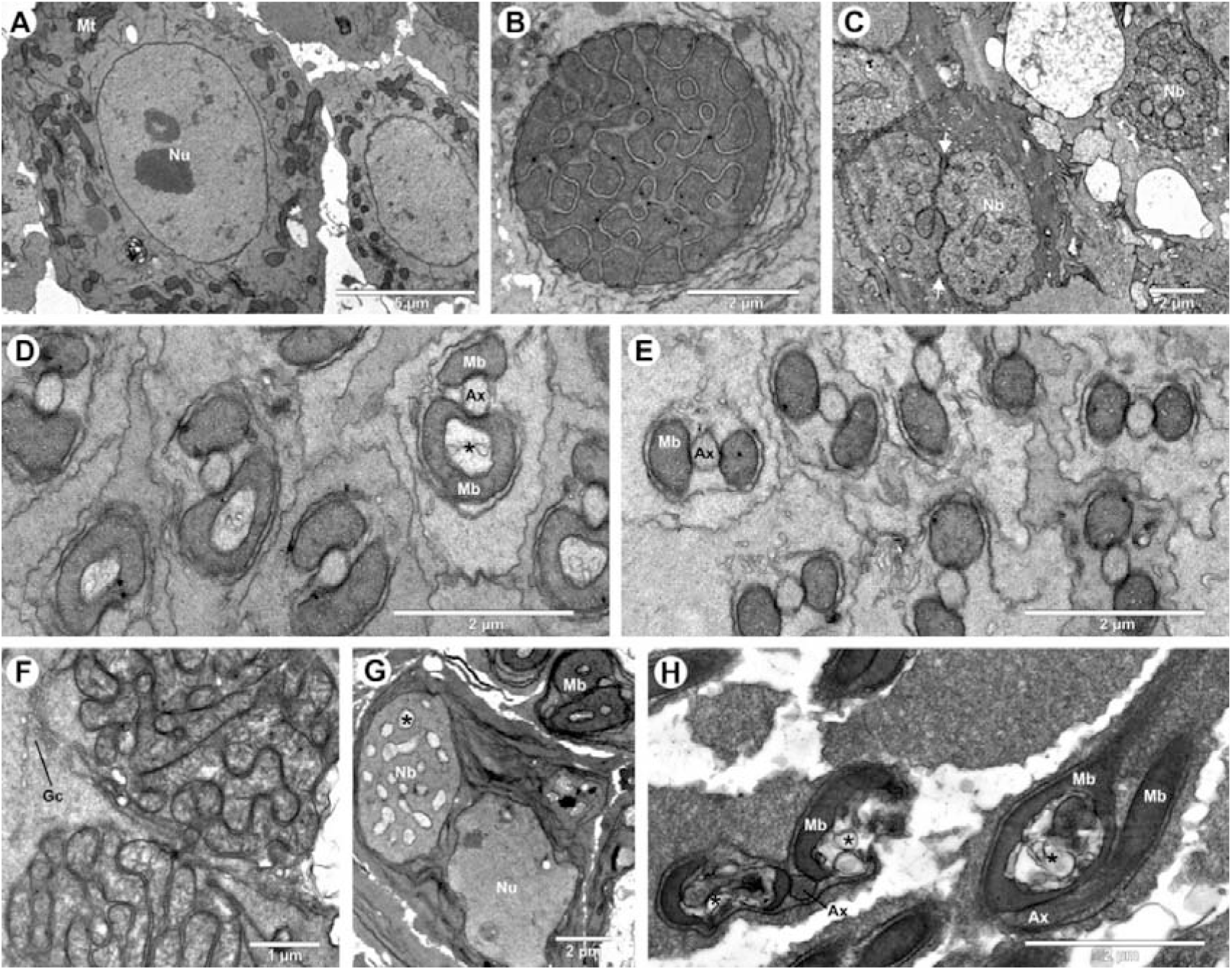
Transmission electron microscopy (TEM) micrographs of mitochondrial development in spermatocytes of non-irradiated (A-E) and irradiated (F-H) *Nezara viridula*. A. Beginning of spermatogenesis, mitochondria cluster. B. Mitochondria fuse into a single nebenkern. C. While further fusing, late nebenkern divide (arrows) into two mitochondrial derivatives. D. Mitochondrial derivatives straddle the axoneme of the developing flagella (sometimes temporarily developing inclusions, ^∗^). E. Late6 stage mitochondrial derivatives. F. Two nebenkerns. G. Typical appearance of nebenkern in irradiated bugs with vacuole-like inclusions. H. Vacuoles (^∗^) and distortion of late stage flagella in irradiated bug. Key: Ax, axoneme; Gc, Golgi complex; Mb, mitochondrial derivatives; Mt, mitochondria; Nb, nebenkern; Nu, nucleus.

Nuclear and sperm head development were also affected by radiation, in particular the organisation of the base of the developing flagella. In both non-irradiated (Fig. 3A) and irradiated insects (Fig. 3B), the gross pattern of sperm development was generally similar. Inside the nucleus there were various structures (e.g., electrondense patches and fibrils) and arrangement of nuclear and cytoplasmic structures developed to give the cell a clear anterior and posterior. Tail development was frequently disrupted in the irradiated insects particularly the microtubule organising centre (MTOC) and accessory machinery. In non-irradiated *N*. *viridula*, we observed a single MTOC attached to the nuclear membrane, which was surrounded by centriole adjuncts, a dense material comprised fused granules (Fig. 3C). In irradiated insects, there was significant variation in morphology at the posterior of the nucleus. Cells were observed with a single MTOC with adjuncts (normal), but also cells were observed with single MTOC but no adjuncts, and cells with two MTOCs (Fig. 3D), either with or without adjuncts.

**Figure 3.**
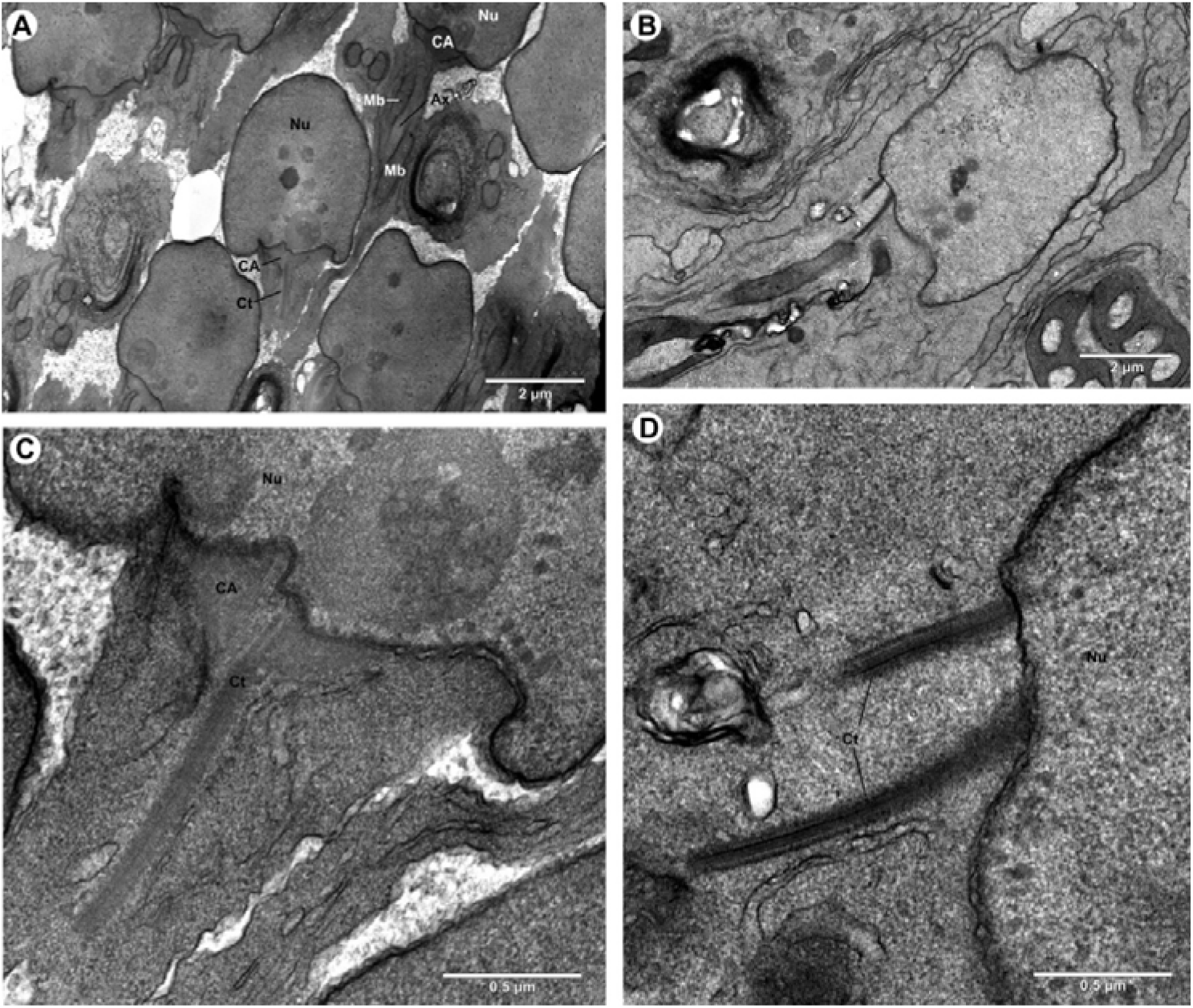
Transmission electron microscopy (TEM) micrographs of early sperm head development in non-irradiated (A, C) and irradiated (B, D) *Nezara viridula*. A. Longitudinal view of early normal sperm development. B. Similar stage in irradiated bug. C. Close up of Figure 3A, microtubule organising centre (MTOC). D. Close up of Figure 3B, two MTOCs in an irradiated bug. Key: CA, centriole adjunct; Ct, MTOC, Mb, mitochondrial-derived body; Nu, nucleus.

All flagella observed in non-irradiated insects maintained an organised morphology during development (Fig. 2D, E), with a single central axoneme surrounded by a pair of mitochondrial derivatives. The fine structure of the axoneme was surrounded by intracellular membrane structures possibly associated with the Golgi complex. In irradiated insects, the morphology was variable (Fig. 4A). Multiple, what appeared to be axonemes, typically two to four, were common and sometimes more than two mitochondrial derivatives were observed. While most observations of multiple axonemes were of discrete structures, occasionally there were either bends or links joining axonemes of uncertain structure observed (Fig. 4B).

**Figure 4.**
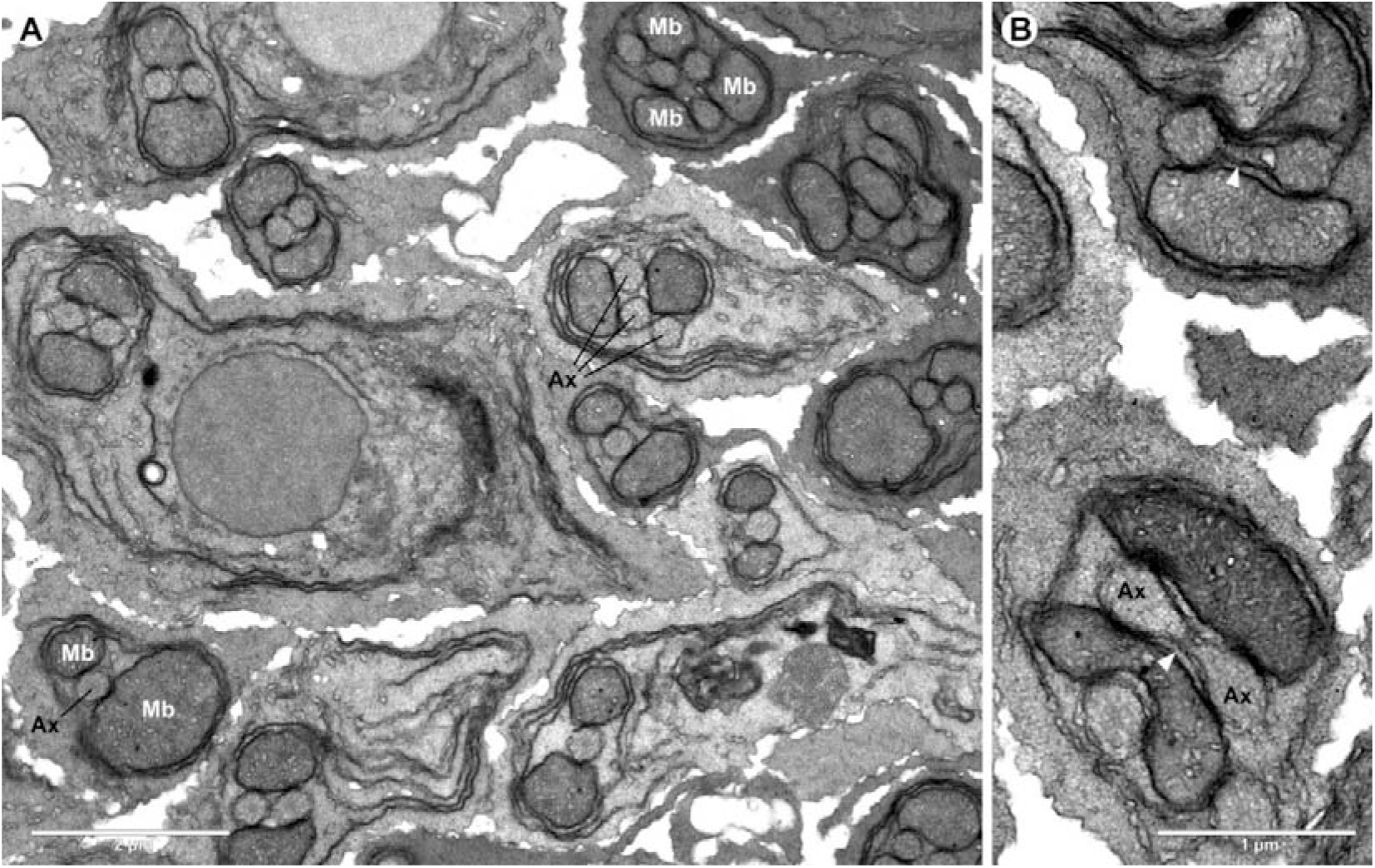
Transmission electron microscopy (TEM) micrographs of variable morphology in the developing tails of irradiated *Nezara viridula*. A. Developing flagella varied in number of axonemes and mitochondrial-derived bodies. B. Connection between axonemes (arrow heads). Key: Ax, axoneme; Mb mitochondrial derivatives.

### 2. Cytochemistry and TEM

Both acid phosphatase and glucose-6-phosphate cytochemical stain procedures selectively enhanced specific features of the testes ultrastructure, rendering those features darker or grainier than the equivalent structures in the control samples which were stained using conventional TEM heavy metal stains (Fig. 5A). Because the efficacy of both acid phosphatase and glucose-6-phosphatase stains were patchy we focused on comparison of well-defined structures, in particular the later stages of flagella development.

**Figure 5.**
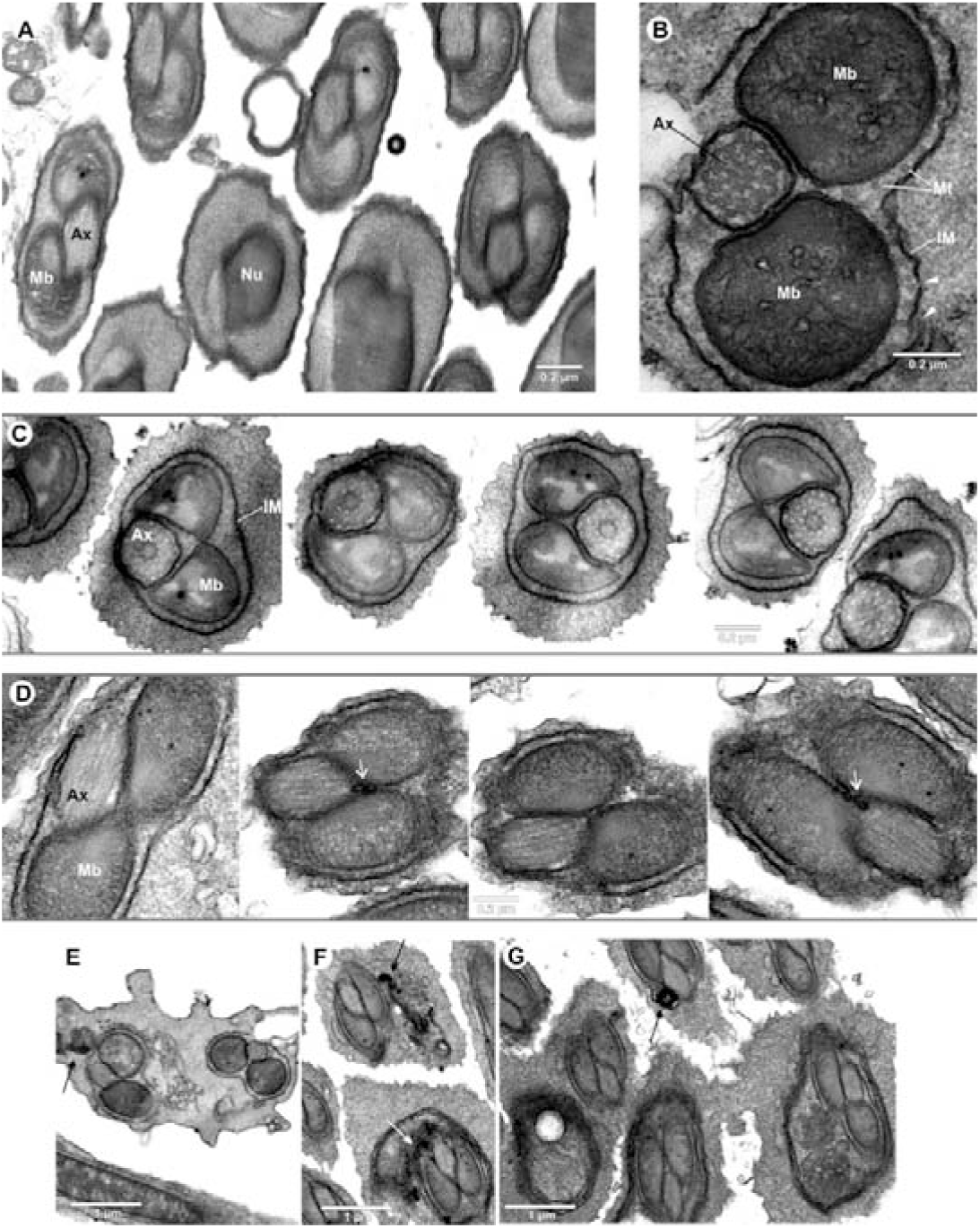
TEM micrographs of late-stage *Nezara viridula* sperm tails in transverse section, indicating the presence of acid phosphatase (AP). A. Non-irradiated sample shows the normal structure of late-stage flagella development with two mitochondrial derived bodies (Mb), which contain electron-dense spots, flanking an axoneme (Ax) and each surrounded by a membrane. Nu, part of the nuclear complex. B. AP non-irradiated sample showing dark specific staining of the discontinuous internal membrane (holes, arrow heads) surrounding the flagella and the continuous membranes around axoneme and mitochondrial derivatives. Mt, microtubules. C. Examples of flagella sections to illustrate staining consistency. D. Irradiated sample with AP staining in membranes, but also in dark lumps in between the mitochondrial derived bodies (Arrows) in some sections. E. Non-irradiated AP sample showing dark stained membranous inclusion adjacent to developing flagella (arrow). F. Similar dark-staining inclusion (arrow) in irradiated sample. G. Inclusion (arrow) in irradiated sample and large dark-stained Golgi complex surrounding flagella at bottom of micrograph and adjacent to flagella (bottom left).

In developing flagella, acid phosphatase appeared to be associated with a discontinuous internal membrane, possibly a modified Golgi complex which surrounded the developing flagella, and also with the membrane which surrounded axonemes and mitochondrial derived bodies (Fig. 5B). The label was consistently located in both non-irradiated (Fig. 5C) and irradiated (Fig. 5D) samples. The main difference was the occasional presence of heavily stained granules in the irradiated sample in between the mitochondrial derived bodies, which were not observed in the non-irradiated sample (Fig. 5D). Heavily stained structures of uncertain globular appearance, which may have been composed of multiple closely apposed membranes, were observed close to the developing flagella in both non-irradiated (Fig. 5E) and irradiated samples (Fig. 5F, 5G).

In the wider population of cells, similar heavily stained globular structures were occasionally observed in both irradiated and non-irradiated samples and appeared to be a modification of, or at least often associated with, Golgi complex. The Golgi complex was often but not always darkly labelled with the distribution of staining being similar in both non-irradiated (Fig. 6A) and irradiated (Fig. 6B) samples. In both samples the nuclear membrane was also typically darkly labelled.

**Figure 6.**
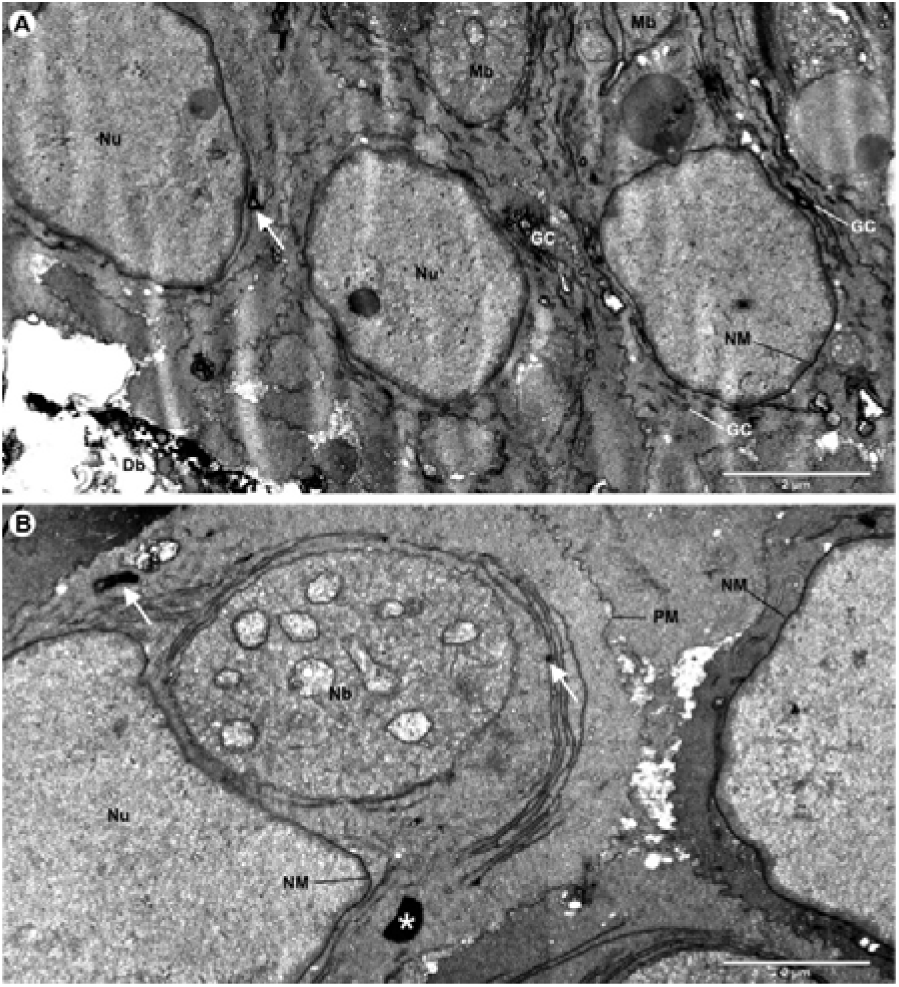
TEM micrographs showing acid phosphatase labelling in early-stage developing sperm cells of *Nezara viridula*. A. Some, but not all, Golgi complex (GC) was well labelled along with nuclear membranes (NM). Extracellular debris (Db) was often heavily labelled. Darkly stained inclusions (arrow) of uncertain identity were also observed sporadically (Arrow). B. Similar observations in irradiated sample, with some spots especially heavily stained (arrow) and some probable artefacts (^∗^). Key: Nu, nucleus; Mb, mitochondrial derived body; Nb, nebenkern (consolidated mitochondria).

In the late-stage developing sperm flagella observations of glucose-6-phosphatase, cytochemical staining was similar to acid phosphatase except that the membrane surrounding the individual mitochondrial derivatives were more heavily stained, while the membrane surrounding the axoneme and the discontinuous membrane surrounding the whole flagella were less heavily stained in both the non-irradiated (Fig. 7A) and irradiated (Fig. 7B) samples.

**Figure 7.**
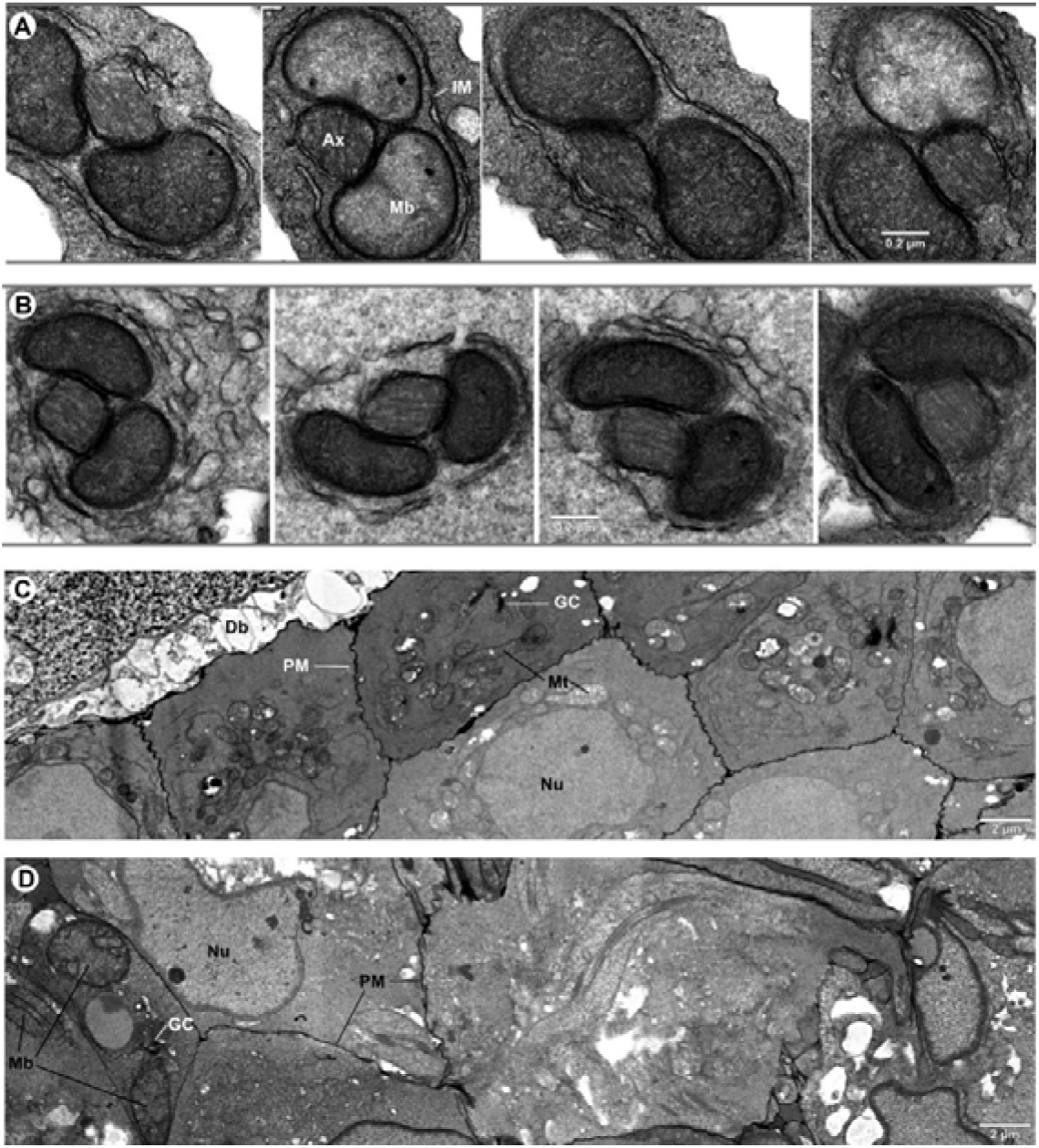
TEM micrographs showing glucose-6-phosphatase labelling in developing *Nezara viridula* sperm cells. A. Transverse view of developing flagella showing lightly labelled internal discontinuous membrane (IM) and axoneme (Ax) membrane and heavily stained membrane surrounding mitochondrial derivatives (Mb). B. Similar staining pattern observed in irradiated sample. C. Early stage sperm cells showing labelling in extracellular debris (Db), the plasma membranes (PM) and in some part of the Golgi complex (GC). No nuclear (Nu) staining was observed.

In cells at a wide range of different stages of sperm development, we observed that staining for glucose-6-phosphatase was less prominent in Golgi than was staining for acid phosphatase, appearing to stain only some complexes or some parts of some complexes. Unlike acid phosphatase, glucose-6-phosphatase staining was intense within cell plasma membranes, but there was no staining of nuclear membranes (Fig 7C, 7D). Although it is possible that fewer Golgi complexes were stained for glucose-6-phosphate in the irradiated (Fig. 7D) compared to the non-irradiated sample (Fig. 7C), this could not be quantified. The glucose-6-phosphatase irradiated sample often contained tissue which was highly disorganised, with cells at multiple stages of development often found together and all oriented in different directions, something not observed in the non-irradiated samples nor the irradiated acid-phosphatase or equivalent control samples.

### 3. Chromosome Complement

Non-irradiated testes showed the expected 2n = 14 - 6 bivalents plus X and Y (Fig 8A) at metaphase 1. There was usually five rod and one ring bivalent (90%) of cells, although 7% showed six rod bivalents and in 2% of cells there were two ring and four rod bivalents. Among the autosomal chromosomes there is one large bivalent (designated A1 by Camacho et al. 1985) and five smaller ones.

**Figure 8.**
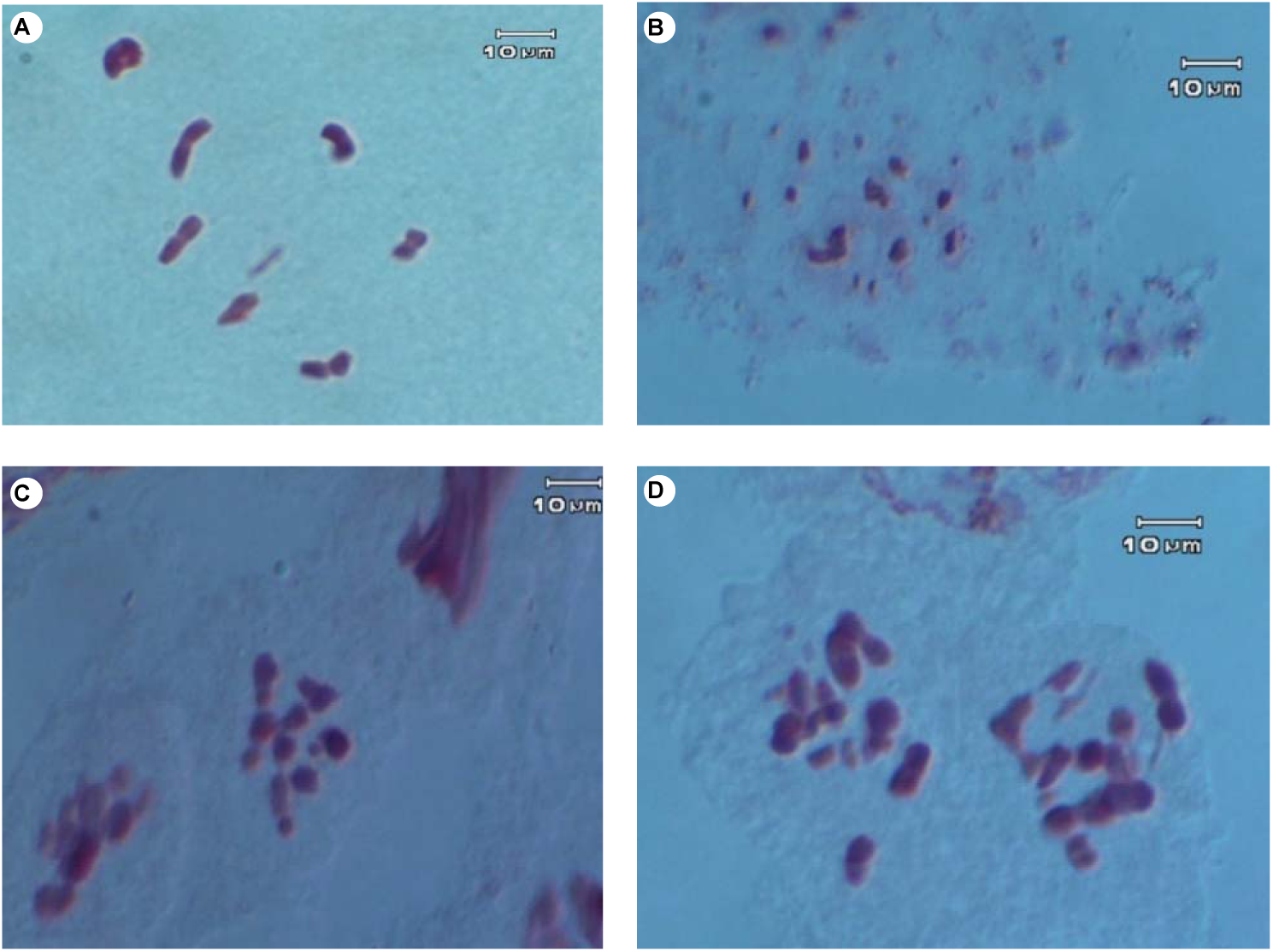
Chromosomes of *Nezara viridula* at metaphase I showing: A. Non-irradiated, normal complement of chromosomes – 6 autosomes and XY, B, C, D: fragmentation of chromosomes and formation of chromosome chains in testes from males irradiated with 40Gy

The testes from four insects that were irradiated with 40Gy all showed fragmented chromosomes at metaphase 1 of meiosis with a range of severity for each insect. A sample of these is shown in figure 8B where the chromosomes are very fragmented, and Figures 8C and 8D where the chromosome are less severely fragmented. No normal divisions were observed. At anaphase II lagging chromosomes, and abnormal divisions (e.g. with 3 spermatids) were observed.

## Discussion

Transmission electron microscopy ultrastructure of the spermatids and spermatozoon in non-irradiated *N*. *viridula* has been described in detail by Fernandes and Báo (1998). The spermatids undergo a series of modifications resulting in the formation of a highly differentiated cell, the spermatozoon which consists of a head containing the nucleus and acrosome, and the tail which is formed by axoneme and two mitochondrial derivatives. To evaluate the impact of irradiation dose on spermatogenesis we selected a dose of 40Gy.

We observed morphologically significant variation in the small number of samples we investigated. However, not all the irradiated samples had extreme differences in morphological features, such as supernumerary axonemes and mitochondrial derivatives, instead only having a few abnormalities while the rest of the structures appeared to be comparable to control insects. Ameresekere et al. (1971) described morphological and histological changes resulting from exposure of nymphs and adults of the hemipteran leaf hopper *Circulifer tenellus* to ^60^Cobalt radiation. Treatment of those males resulted in pronounced cell necrosis and sperm depletion, as well as reduction in various stages of spermatogenesis and vacuolation of the testicular sacs, with slight thickenings of the walls. It has been reported that X-ray dosages of 2.5, 5.5 and 8.5 kR (≈ 22, 48 and 74 Gy) caused degeneration and necrosis in reproductive organs, which resulted in empty cysts, in the house fly, *Musca domestica* (L.); in the secondary screwworm fly, *Cochliomyia macellaria* (Fab.); and in the black blow fly, *Phormia regina* (Meigen) (Riemann 1967, Riemann and Thorson 1969). Banu et al. (2006) had reported that the gamma radiation effect on the gonads of red flour beetle *Tribolium castaneum* treated with 15 Gy resulted in significant size reduction of testes and ovaries, leading to sterility in both sexes. A significant reduction in the size and functionality of *N*. *viridula* reproductive organs would probably lead to reduction in overall fecundity and fertility. For a sterile insect release programme to be successful, irradiated insects need remain sexually competitive (Knipling 1955). Any reduction in fitness would require a larger number of insects to be released (Kean et al. 2011).

All mitochondrial stages are very sensitive to radiation (Coggins 1973). We found that the nebenkern did not form normally in irradiated individuals. These findings were similar to (Coggins 1973), who theorised that the failure of the mitochondria to fuse into a nebenkern may due to temporary interference with either the genes controlling this formation process and/or the breakdown or the formation of the bounding membrane.

We observed supernumerary and abnormal structures in the irradiated insects. Similar results have been observed in irradiated red palm weevils *Rhynchophorus ferrugineus* with multiple axonemes (Paoli et al. 2014), also in irradiated desert locusts *Schistocerca gregaria*, where supernumerary centrioles, mitochondrial material, flagella and nuclear outgrowths have been described (Gatenby 1941, Coggins 1973). Tahmisian and Devine (1961) demonstrated the presence of supernumerary acrosomes and centrioles around the spermatid nuclei of irradiated *Melanoplus differentialis* grasshopper. We observed non-symmetrical mitochondrial derivatives in the irradiated insects, unlike what we observed in our non-irradiated insects, where mitochondrial derivatives appeared symmetrical, as is expected in non-irradiated Pentatomidae (Araújo et al. 2011). The axoneme and mitochondrial derivative bodies provide motility to the sperm; abnormalities here would probably reduce the ability of the sperm to swim via the whipping of the flagellum. The axoneme gives structure to the flagellum, yet is flexible. An excess of axonemes may increase the rigidity of the flagellum, reducing the amount of thrust available to the sperm or produce multiple-tailed sperm (du Plessis and Soley 2011). Abnormalities in the mitochondrial derivatives in sperm could reduce its likelihood of reaching any ova (Coggins 1973, Paoli et al. 2014).

The investigation in chromosome structure mimicked the results from the TEM and testes visual structure, in that individual insects were affected differently even though they were all exposed to the same dose of radiation. In some irradiated testes, the chromosomes were extremely fragmented while in others there was much less fragmentation and some cells appeared similar to the non-irradiated. In this study we were unable to obtain any progeny from any males that had been irradiated with 40Gy as the males showed low physical activity and did not mate.

The ultrastructure of the spermatids and the sperm of *N*. *viridula* demonstrated a severe effect of gamma radiation on the gonads and spermatogenesis. Further replication over a variety of doses coupled with biological studies would help to quantify the variation in spermatogenesis abnormalities between individuals exposed to radiation, as well as sterility rates. In terms of the SIT technique, a requirement is that individuals are able to mate successfully and maintain sterility in the population, so a dose lower than 40Gy would need to be administered to any insects to maintain physical fitness. The potential for inherited sterility in stink bugs (Pentatomidae) requires further investigation particularly at the lower levels of irradiation.

## Acknowledgments

This work was funded by Plant & Food Research through the Better Border Biosecurity collaboration (www.b3nz.org), and by AgResearch Protein and Biomaterials team. We wish to thank James Vernon, Santanu Deb Choudhury and Hatem Mohamed for technical assistance and advice, and Paul Sutherland and Kye Chung Park for reviewing earlier versions of the manuscript.

